# Improved cognitive-motor processing speed and decreased functional connectivity after high intensity aerobic exercise in individuals with chronic stroke

**DOI:** 10.1101/2023.01.15.523513

**Authors:** Justin W. Andrushko, Shie Rinat, Brian Greeley, Beverley C. Larssen, Christina B. Jones, Cristina Rubino, Ronan Denyer, Jennifer Ferris, Kristin L. Campbell, Jason L. Neva, Lara A. Boyd

**Affiliations:** Brain Behaviour Laboratory, Vancouver British Columbia, Canada, V6T 1Z3; Faculty of Medicine, Department of Physical Therapy, University of British Columbia – Vancouver, British Columbia, Canada, V6T 1Z3; Faculty of Medicine, Graduate Program in Rehabilitation Sciences, University of British Columbia, Vancouver, British Columbia, Canada, V6T 1Z3; Faculty of Medicine, Graduate Program in Neuroscience, University of British Columbia, Vancouver, British Columbia, Canada, V6T 1Z3; Faculty of Medicine, School of Kinesiology and Physical Activity Sciences, University of Montreal, Montreal, Quebec, Canada, H3T 1J4; Research Center of the Montreal Geriatrics Institute (CRIUGM), Montreal, QC, Canada

**Keywords:** Chronic stroke, functional connectivity, high intensity aerobic exercise, trail making test, cognitive function

## Abstract

After stroke, impaired motor performance is linked to an increased demand for cognitive resources. Aerobic exercise improves cognitive function in healthy populations and may be effective in altering cognitive function post-stroke. We sought to determine if high intensity aerobic exercise paired with motor training in individuals with chronic stroke alters cognitive-motor function and functional connectivity between the dorsolateral prefrontal cortex (DLPFC), a key region for cognitive-motor processes, and the sensorimotor network. Twenty-five participants with chronic stroke were randomly assigned to exercise (n = 14; 66 ± 11 years; 4 females), or control (n = 11; 68 ± 8 years; 2 females) groups. Both groups performed five-days of paretic upper limb motor training after either high intensity aerobic exercise (3 intervals of 3 minutes each, total exercise duration of 23-minutes) or watching a documentary (control). Resting-state fMRI, and TMT-A and B were recorded pre- and post-intervention. Both groups showed implicit motor sequence learning (*p* < .001), but there was no added benefit of exercise (*p* = .738). Regardless of group, the changes in task score (*p* = .025), and dwell time (*p* = .043) were correlated with a decrease in DLPFC-sensorimotor network functional connectivity (*p* = .024), which is thought to reflect a reduction in the cognitive demand and increased automaticity. The exercise group experienced greater overall cognitive-motor improvements measured with the trail making test part A (TMT-A: task score: *p* = .012; dwell time: *p* = .024; movement time: *p* = .567). Aerobic exercise may improve cognitive-motor processing speed post-stroke.

**Significance statement:** After stroke, impaired motor performance is linked to an increased demand for cognitive resources. In our work we show that high intensity aerobic exercise paired with an implicit motor learning task improves cognitive-motor processing speed and reduces resting-state functional connectivity between the dorsolateral prefrontal cortex and the sensorimotor network in individuals living with chronic stroke. These data likely reflect a reduction in cognitive resource dependence during a cognitive-motor task after stroke and a shift towards cognitive-motor automaticity.

## Introduction

Roughly 15 million people experience a stroke each year (Mittmann *et al*., 2012). Stroke is the second most common cause of death globally and one of the leading causes of severe, adult disability (Katan & Luft, 2018). Due to advancements in preventive care, rates of stroke declined between 1990 and 2016, yet the number of individuals that survive and live with severe disability nearly doubled during that same timeframe (Lindsay *et al*., 2019). Identifying methods to enhance recovery from stroke to improve or maintain independence of living is an important and persistent research objective.

After stroke, cognitive impairment may interact with or influence motor recovery. Greater cognitive resources are needed to successfully plan and execute voluntary movements after stroke (Puh *et al*., 2007). This impaired cognitive demand is observed through an increase in cortical activity in prefrontal areas including the dorsolateral prefrontal cortex (DLPFC) (Puh *et al*., 2007; Meehan *et al*., 2011; Li *et al*., 2014), and is often associated with worse motor function (Meehan *et al*., 2011; Li *et al*., 2014; Lin *et al*., 2021; Hall *et al*., 2021). The DLPFC is an important brain region involved in several cognitive-motor processes (i.e., cognitive processes involved in cognitively demanding motor tasks) including processing speed (Hillary *et al*., 2006; Kaller *et al*., 2011), response selection (Boyd *et al*., 2009*b*), and task switching (Badre & Wagner, 2004; Hart *et al*., 2013; Brunoni & Vanderhasselt, 2014). Importantly, past work from our lab showed that despite an equal dose of motor practice and subsequent learning, individuals with chronic stroke were unable to shift cortical activity away from the prefrontal cortex during a motor learning task (rather, an increase was observed), while age matched healthy controls did (Meehan *et al*., 2011). This shift in cognitive resources away from the DLPFC during cognitive-motor tasks has been observed in healthy cohorts with enhanced automaticity coinciding with a decrease in functional connectivity between the DLPFC and the sensorimotor network (Mazzoni, 2008; Wu *et al*., 2008). Therefore, interventions that can reduce the cognitive demand and DLPFC activity in individuals with stroke may improve motor performance.

Aerobic exercise can alter patterns of brain activity, including resting-state functional connectivity between brain networks as measured by functional magnetic resonance imaging (fMRI) (Weng *et al*., 2017; Greeley *et al*., 2021), and is known to improve cognitive function in healthy older adults (Barnes, 2015), and in individuals with stroke (Zheng *et al*., 2016). High intensity aerobic exercise in particular can also improve motor skill acquisition in neurologically intact people (Skriver *et al*., 2014; Stavrinos & Coxon, 2017; Dal Maso *et al*., 2018; Kendall *et al*., 2020), and individuals with chronic stroke (Nepveu *et al*., 2017). These improvements are linked to a decrease in the neurotransmitter gamma-aminobutyric acid (Singh *et al*., 2014; Singh & Staines, 2015; Stavrinos & Coxon, 2017; Hendy *et al*., 2022), and an increase in the protein brain-derived neurotrophic factor (Sleiman *et al*., 2016), both of which play important roles in neuroplasticity (Stagg *et al*., 2011; Mang *et al*., 2013; Andreska *et al*., 2020). Therefore, beyond the potential cognitive benefits associated with high intensity aerobic exercise, pairing it with motor rehabilitation is a promising method to enhance cognitive-motor function.

The purpose of the current study was to determine if high intensity aerobic exercise paired with skilled motor training alters cognitive-motor function and DLPFC-sensorimotor network functional connectivity in individuals living with chronic stroke. To address these questions participants were tested using the cognitive-motor assessments Trail Making Test parts A (TMT-A) and B (TMT-B). The Trail Making Tests are commonly used to assess executive function and can measure processing speed with TMT-A, and mental flexibility and task switching with TMT-B (Kortte *et al*., 2002; Bowie & Harvey, 2006; Gläscher *et al*., 2012; MacPherson *et al*., 2017). Additionally, resting-state fMRI scans were acquired to measure change in functional connectivity. We hypothesized that high intensity aerobic exercise paired with skilled motor training would enhance cognitive-motor performance as measured with TMT-A and TMT-B in individuals living with chronic stroke, and these changes would be correlated with a decrease in DLPFC-sensorimotor network functional connectivity, reflecting a reduction in cognitive resources needed to perform the tasks.

## Methods

### Ethical approval

This study conformed to the standards set by the Declaration of Helsinki and was approved by the University of British Columbia Clinical Research Ethics Board: #H16-01945.

### Participants

Individuals living with chronic stroke (> 6 months post-stroke) were recruited to participate in this study if they had an ischemic or hemorrhagic stroke and were between the ages of 21-85 years old. In addition, to participate individuals had to score > 23 on the Montréal Cognitive Assessment (MoCA) (Nasreddine *et al*., 2005), and be cleared by a cardiologist for safe participation in an exercise protocol after performing a supervised stress-test. Eligible participants were randomly allocated to either an exercise or control group. Both groups performed the same motor training intervention immediately following either aerobic exercise or rest. The data in this manuscript come from a large study on the impact of exercise on behaviour, brain function, and physiology in individuals with stroke. The data reported here are a subset of the larger study (Greeley *et al*., 2021, 2023; Neva *et al*., 2022).

### Experimental design

After obtaining informed and written consent, participants performed a graded maximal exercise stress test to determine their eligibility to participate in the intervention aspects of this study. Testing was done before (pre-testing) and after (24h-post) five-days of skilled motor training using the paretic arm paired with either high intensity aerobic exercise (exercise group), or rest (control group). Resting-state fMRI, TMT-A, TMT-B, Fugl-Meyer Assessment of Motor Recovery (FMA) (Fugl Meyer *et al*., 1975), and the Wolf Motor Function Test (WMFT) (Wolf *et al*., 2001) were completed at pre-testing and 24h-post testing time points to determine brain - behaviour relationships (Figure 1).

**Figure 1.**
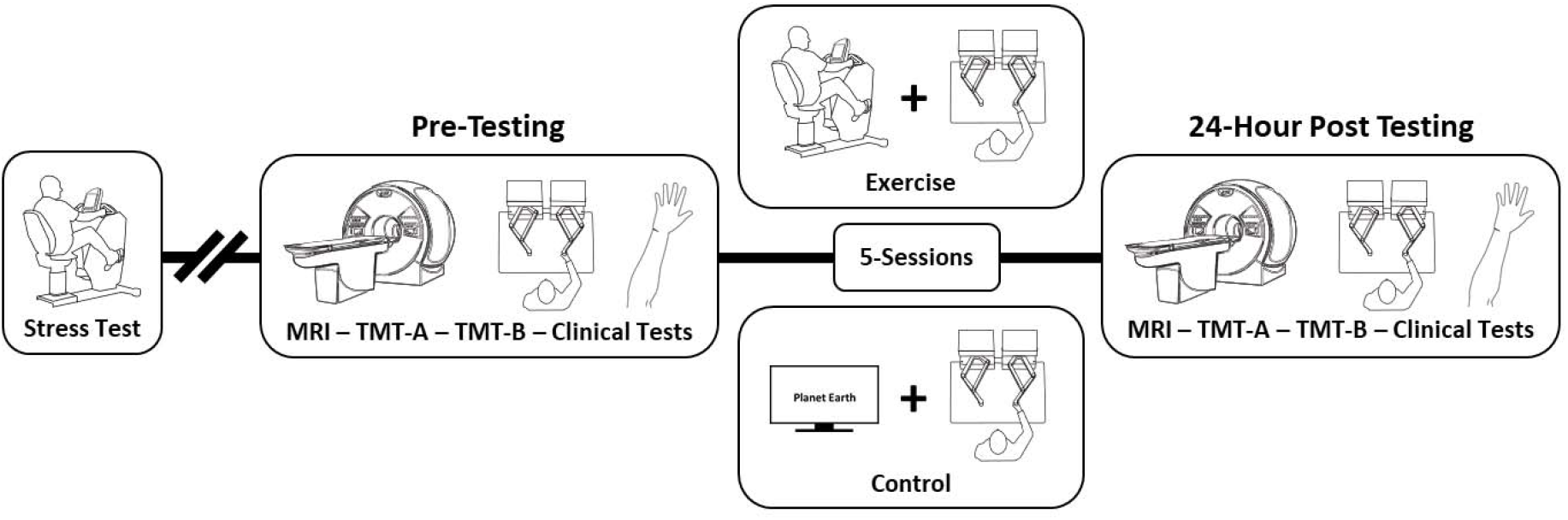
Experimental design timeline. MRI = magnetic resonance imaging, TMT-A & B = Trail Making Test Part A and B, Clinical tests = Fugl-Meyer Assessment, and Wolf Motor Function Test at pre-testing and 24h-post timepoints, and Montréal cognitive assessment at pre-testing only.

### Stress test

The graded maximal exercise stress test was administered by a cardiology technician. Electrocardiogram electrodes were used for continuous heart rate monitoring throughout the exercise protocol. Prior to the test participant lay supine for approximately three minutes, after which resting heart rate (HR), blood pressure (BP) and rating of perceived exertion (RPE) were recorded. Next, the participant was seated in an upright recumbent bike (SCIFIT, Tulsa, Oklahoma, USA), that was adjusted to fit the participant. During the stress test, HR and RPE were recorded every minute and BP was recorded every two minutes. Participants were instructed to maintain a cadence between 50-80 revolutions per minute (RPM) and that dropping below 50 RPM would terminate the test. The stress test began with a two-minute warm-up at 10 Watts (W) of resistance. Following the warm up, the wattage was increased by 5, 10, or 15 W depending on subjective observation of performance (Beltz *et al*., 2016). The resistance was increased every minute until the participant was unable to maintain a 50 or greater RPM cadence or when volitional fatigue was reached. Once the participant reached the termination criteria, the resistance was dropped back to 10 W for approximately three minutes as a cool down. Next, the participant moved back to the supinated resting position until their HR, BP, and RPE recovered back to baseline levels.

### Groups

#### High intensity aerobic exercise

Each exercise session was completed on an upright recumbent bike (SCIFIT, Tulsa, Oklahoma, USA). Each session started with a five-minute warm-up at 10 watts. Following the warm-up, the participants performed three, three-minute intervals of cycling at 75% of their maximum power output, based on the maximum power output achieved during the final fully completed minute during the exercise stress test. Each interval was separated by three-minutes of low intensity cycling against 10 watts of resistance. The total duration of each aerobic exercise session lasted 23-minutes. BP, HR, and RPE were recorded every three-minutes until the end of the protocol. Immediately following the exercise protocol, participants proceeded with motor training.

#### Control

Participants allocated to the control group watched a Planet Earth documentary for 23-minutes immediately prior to engaging in motor training each session. Heart rate was recorded every three-minutes throughout the video.

### Motor training

The serial targeting task was employed as the motor training task (Brodie *et al*., 2014; Mang *et al*., 2016; Greeley *et al*., 2021). Participants used their paretic arm to control a frictionless manipulandum to move a cursor between a start position and an end target projected by the Kinarm end-point robot (Kinarm, Kingston, ON, Canada). Targets appeared one at a time; as soon as participants finished movement to one target, they were required to hold that positioning for 500 ms for the next appeared. Participants had 10,000 ms to reach each target. A repeated six-element sequence of movements was embedded between seven-element random sequences. The inclusion of both sequences allows us to separate improvements in motor learning (repeated sequence) from those associated with motor control (random sequences) (Boyd *et al*., 2009*a*). In each of the five-training-sessions, participants practiced four-blocks of the serial targeting task (444 movements per session).

Data were analyzed using an exponential curve-fitting algorithm that enables parameterization of motor data across practice (Wadden *et al*., 2017, 2019). Motor learning related change was characterized by fitting behavioural data to an exponential equation:

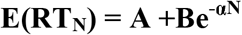

where **RT** is reaction time, **A** is predicted asymptote in performance, **B** is the performance change score to predicted asymptote, **α** is rate of change and **N** is number of practice trials (Brown & Heathcote, 2003). Our dependent measure of motor learning was a change score (**B**) extracted from individual learning curves for each individual’s practice sessions by group (exercise, control).

### Cognitive-motor testing

To assess cognitive-motor performance TMT-A and TMT-B were performed by the participants with their less-affected arm, or in scenarios where there were bilateral lesions participants were instructed to use their preferred arm on a Kinarm end-point robot. The use of the less affected limb allowed us to assess cognitive performance without stroke related motor impairment affecting responses. TMT-A involves connecting dispersed numbered targets in ascending numerical order from ‘1’ to ‘25’ with linear reaching movements. The objective is to locate the next number and reach to it without touching any other numbers or connected lines from the already completed reaches. This test assesses cognitive-motor processing speed, in which the participant must visually navigate, identify the appropriate target, and execute a movement as fast and accurately as possible (Corrigan & Hinkeldey, 1987; Bowie & Harvey, 2006).

TMT-B is similar to TMT-A in that it also involves searching and connecting targets in ascending order. However, TMT-B differs in that the task must be completed in an alternating numeric and alphabetic sequence, where the number ‘1’ must connect to the letter ‘A’, then ‘A’ must connect to the number ‘2’, until the final target number ‘13’ is reached, still equaling 25 total targets. The added complexity stresses the cognitive system and requires mental flexibility and task switching to complete. For TMT A and B, different sequences were used at pre-testing and 24h-post to avoid any possible learning effects.

Three metrics of task performance were assessed for TMT-A and TMT-B. First, to assess overall performance the task score at each time point was used. This metric provides a global measure of an individual’s performance. Specifically, the task score measures deviations from an individual’s best performance. Task scores are always positive with zero representing best performance and deviations from zero reflecting poorer performance (See Dexterit-E Explorer manual for more information - https://kinarm.com/support/user-guides-documentation/). Additionally, to separate the cognitive and the motor aspects of these assessments the total dwell time (i.e., the time that the participant remains on a target while they visually search and plan their next movement) was subtracted from the total task time to isolate the movement time. Then statistical analyses were carried out on the movement time and dwell time separately to determine the impact of the intervention on the cognitive and motor aspects of task performance.

### Clinical assessments

Paretic arm impairment was quantified using the upper extremity portion of the Fugl-Meyer Assessment (FMA; 0-66; higher scores indicate less paretic arm impairment) (Lin *et al*., 2004). The 17-item version of WMFT was used to characterize arm motor function (Wolf *et al*., 2001). The WMFT contains 15 timed movement tasks. For each WMFT task, the rate (repetitions/60 seconds, with a rate of zero recorded if no repetitions were completed within 120 seconds) was calculated to characterize functional impairment (Hodics *et al*., 2012); higher scores reflect a faster movement rate and thus greater motor function. All assessors for FMA and WMFT were trained physical or occupational therapists, or a clinical student in training.

### Magnetic resonance imaging

#### Magnetic resonance imaging acquisition

Participants received structural and functional brain scans on a Philips Achieva 3 tesla or a Philips Elition 3 tesla MRI. At both testing time points a T1-weighted (T1w) structural brain scan (TR = 8.1 ms, TE = 3.61 ms, flip angle = 8°, 1mm^3^ isotropic voxels, field of view = 256 × 256 × 165mm field of view, total scan time = 6.4 minutes), and a resting-state fMRI scan (TR = 2.000 ms, TE = 30 ms, flip angle = 90°, 120 volumes, voxel dimensions = 3 × 3 × 3 mm with a 1 mm gap, total scan time = 4 minutes) were acquired. During resting-state fMRI scans participants were asked to look at a fixation cross, to think of nothing and stay awake.

#### Anatomical data preprocessing

Anatomical T1w images were preprocessed using the fMRIprep pipeline (v22.0.0) (Esteban *et al*., 2018). Briefly, for each participant, the two T1w images from pre-testing and 24h-post were first corrected for intensity inhomogeneity using N4BiasFieldCorrection (Tustison *et al*., 2010) as part of ANTs (v2.3.1). Next, both T1w images were used to create a participant specific average template using mri_robust_template (Reuter *et al*., 2010) from FreeSurfer (v6.0.1), followed by skull-stripping with antsBrainExtraction.sh using OASIS as a target template.

#### Lesion masking

For lesions, a binary mask was created by manually evaluating the T1w images from session one and drawing a mask over the lesioned tissue in 3D space using ITK-Snap (v3.8.0). These binary lesion masks were used by fMRIprep to assist in the registration steps (Figure 2 for lesion mask overlap).

**Figure 2.**
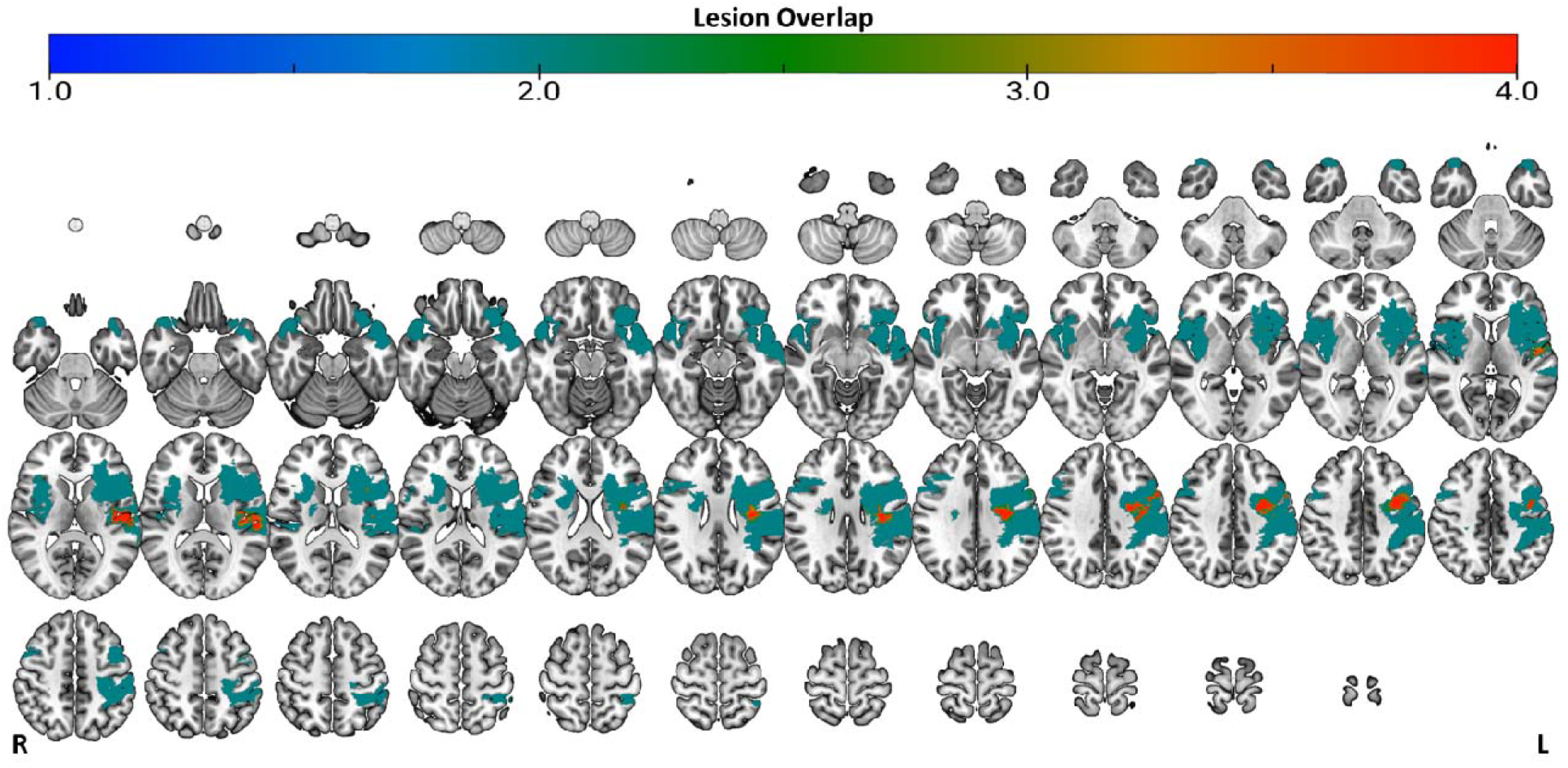
Lesion overlap. Colour bar represents the number of participants that have a lesion in a given location (i.e., lesion voxel overlap; A value of 4 means 4 participants have a lesion in the same location). Figure is in radiological view with the right side of the brain on the left, and the left side of the brain on the right.

#### Resting-state fMRI data preprocessing

Resting-state fMRI data were initially minimally preprocessed in native space using fMRIprep (v22.0.0) (Esteban *et al*., 2018) to carry out fieldmap-less susceptibility distortion correction (Wang *et al*., 2017). Next, MELODIC (v3.15) as part of the FMRIB Software Library (FSL v6.0.3; (Jenkinson *et al*., 2012)) was used to carry out motion correction (MCFLIRT; (Jenkinson *et al*., 2002)), high-pass temporal filtering at 0.01Hz, and the decomposition of the functional runs into independent components for denoising. Following MELODIC preprocessing, FMRIB’s Independent Component Analysis (ICA)-based Xnoiseifier (ICA-FIX) was used to automatically denoise the data (Griffanti *et al*., 2014; Salimi-Khorshidi *et al*., 2014). A custom training-weight was created on a subset of the present study’s data (20 runs total: 10 runs from pre-testing and 10 runs from 24h-post testing) and used with a threshold value of 20 to denoise the data. Finally, after the automated denoising, data were smoothed with a 5 mm full-width half maximum (FWHM) kernel and registered to the MNI152_T1_1mm standard space template included in FSL.

#### Resting-state fMRI data analysis

After preprocessing, a group-level functional connectivity analysis was carried out in FSL’s melodic command-line tool with a dimensionality constraint of 11 group-level components. The constraint of 11 components was selected after multiple cross-correlation analyses between the spatial maps of several different ICA constraints ranging from 5-15 components with the BrainMap 10-ICA template (Smith *et al*., 2009). Dual regression was then performed to estimate a version of the group-level spatial maps for each participant and run (Nickerson *et al*., 2017). The component that best represented the sensorimotor network was then used to constrain a seed-based functional connectivity analysis. A right DLPFC mask was extracted from the Sallet dorsal frontal connectivity-based parcellation (Sallet *et al*., 2013), and used as a seed region for this analysis.

Statistical inference was determined with a four-contrast general linear model (GLM) that compared pre- and post-testing rs-fMRI scans for the exercise group (pre > post, and post > pre), and the control group (pre > post, and post > pre). Non-parametric permutation testing with 5000 permutations for each contrast was carried out using Permutation Analysis of Linear Models (PALM) with family-wise-error-rate corrected contrasts and cluster-extent based thresholding with a z-score of 3.1 (Winkler *et al*., 2014).

### Statistical analysis

To test our hypotheses that high intensity aerobic exercise would enhance less-affected upper limb cognitive-motor performance in individuals living with chronic stroke, as measured with TMT-A and TMT-B, separate mixed repeated measures analysis of variance (RM-ANOVA) tests were performed when data met parametric assumptions. For RM-ANOVA tests, group was entered as a between factor, and time was entered as a within factor variable. When an interaction was significant, a priori pairwise comparisons were performed. For these pairwise comparisons, independent samples *t*-tests, and paired-sampled *t*-tests were used to investigate differences between groups at each time point, and between time points within each group respectively. Additionally, where significant effects were observed, secondary exploratory analyses were performed to assess sex differences with no a-priori hypotheses. Partial eta^2^ (□_p_^2^) effect sizes were reported for all interactions and main effects. Data normality was assessed with Shapiro-Wilk’s tests, and homogeneity of variance was assessed with Levene’s tests.

When data did not meet the appropriate parametric testing assumptions of normality or heterogeneity of variance, Mann-Whitney U tests were used to assess between group differences, and Wilcoxon signed-rank tests were used to analyze paired-sample within group data. Rank-Biserial Correlations were reported as the effect sizes for non-parametric tests.

For all parametric and non-parametric tests, a manual Bonferroni adjusted alpha-level (α = .0125) was then used to reduce family-wise-error rates. To improve clarity in statistical reporting, the uncorrected *p*-values from the individual pairwise tests from the parametric and non-parametric comparisons were Bonferroni adjusted by multiplying the uncorrected *p*-value by the number of comparisons within a given variable (4 tests; 2 within group, and 2 between group comparisons). This is a mathematically equivalent approach to adjusting the alpha-level threshold by dividing it by the number of tests (i.e., α = .0125; α = .05/4 tests), and is the same approach used for Bonferroni post-hoc testing in statistical software packages (https://www.ibm.com/support/pages/calculation-bonferroni-adjusted-p-values). This approach allows significance interpretation to remain at α = .05. Finally, Spearman’s Rho (ρ) correlations were used to test our hypothesis that behavioural changes would relate to a reduction in functional connectivity between the DLPFC and the sensorimotor network. Statistical analyses were performed in JASP (v0.16.4.0) (Love *et al*., 2019).

## Results

### Participants

A total of 41 individuals with stroke consented to participate in this study, however, the exercise stress test revealed abnormalities in three individuals, while two others were ineligible due to low MoCA scores. Eligible participants were randomly assigned to an exercise or control group. Of the remaining 36 individuals, two had missing MRI data, and nine had excessive head motion exceeding a mean framewise displacement greater than .5 mm during MRI scans, thereby rendering at least one of their time points unusable. Therefore, a total of 25 participants were included in this study (Table 1).

**Table 1.** Demographics.

| Group | Age | Sex | Affected hemisphere | Stress test Max watts | MoCA | FMA |  | WMFT |  |
| --- | --- | --- | --- | --- | --- | --- | --- | --- | --- |
|  |  |  |  |  |  | Pre-testing | 24h-post | Pre-testing | 24h-post |
| Exercise<br>n = 14 | 66 ± 11 | F: n = 4<br>M: n = 10 | Left: n = 5<br>Right: n = 9 | 83 ± 32 | 26 ± 2 | 52 ± 16 | 52 ± 16 | 48 ± 27 | 52 ± 36 |
| Control<br>n = 11 | 68 ± 8 | F: n = 2<br>M: n = 9 | Left: n = 6<br>Right: n = 5 | 73 ± 31 | 25 ± 2 | 53 ± 12 | 54 ± 11 | 37 ± 17 | 42 ± 20 |

MoCA = Montréal Cognitive Assessment; FMA = Fugl-Meyer Assessment; WMFT = Wolf Motor Function Test

### Clinical assessments

Separate group × time RM-ANOVA tests for FMA and WMFT were assessed. The ANOVA failed to detect a significant group × time interaction in FMA score [*F*(1, 22) = .286, *p* = .598, □_p_^2^= .013], nor main effects of time [*F*(1, 22) = .750, *p* = .396, □_p_^2^= .033], or group [*F*(1, 22) = .132, *p* = .719, □_p_^2^= .006]. Similarly, a group × time interaction [*F*(1, 20) = .017, *p* = .897, □_p_^2^< .001], and main effects of time [*F*(1, 20) = 3.391, *p* = .080, □ ^2^= .145], and group [*F*(1, 20) = 1.052, *p* = .317, □_p_^2^= .050] were not significant for the mean rate of performance from the WMFT.

### Serial targeting task

For the B values there was a violation of normality for the exercise group as assessed with Shapiro-Wilk’s test (W = .513, *p* < .001). Therefore, a non-parametric Mann-Whitney U test was used to assess between group differences. There was no statistically significant difference between exercise (Mean ± SD; .364 ± .352 B value) and control (.315 ± .122 B value) groups (*W* = 71, *p* = .738, Rank-Biserial Correlation = .092). However, both groups learned the task, and improved their motor performance throughout the intervention, as evidenced by a one-sample Wilcoxon signed-rank test, which indicates that the B values were significantly different from zero (V = 276, *p* < .001, Rank-Biserial Correlation = 1.000; Figure 3).

**Figure 3.**
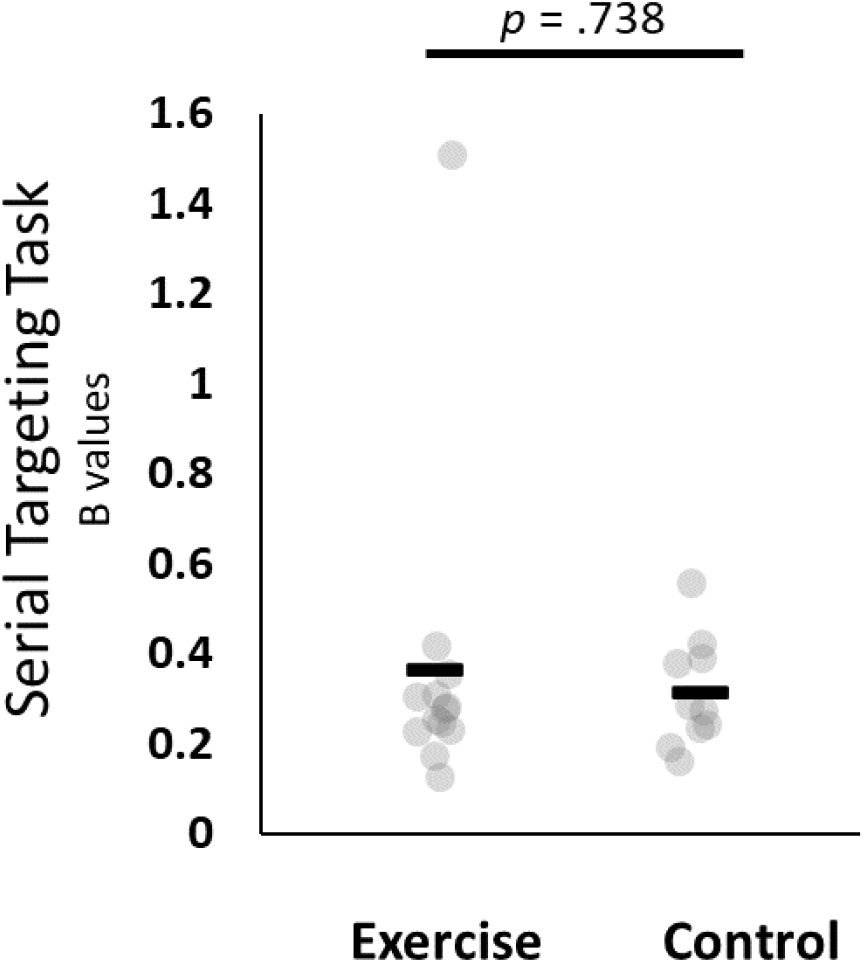
Serial targeting task B values. There were no differences between groups (p = .738), but data were significantly different from zero (p < .001). Black bars represent the group means. Grey circles represent individual data points.

### Trail making test part A

#### Task score

For TMT-A, a RM-ANOVA test revealed a significant group × time interaction [*F*(1, 22) = 9.257, *p* = .006, □_p_^2^= .296], and a significant main effect for time [*F*(1, 22) = 9.269, *p* = .006, □_p_^2^= .296], but the main effect of group was not significant [*F*(1, 22) = .761, *p* = .393, □_p_^2^= .033]. Bonferroni adjusted post-hoc testing revealed that the interaction was influenced by a significant pre- to 24h-post testing difference for the exercise group only (-.673 ± .150, *t* = 4.496, *p* = .012). No other post-hoc tests were statistically significant (all *p* > .180; Figure 4).

**Figure 4.**
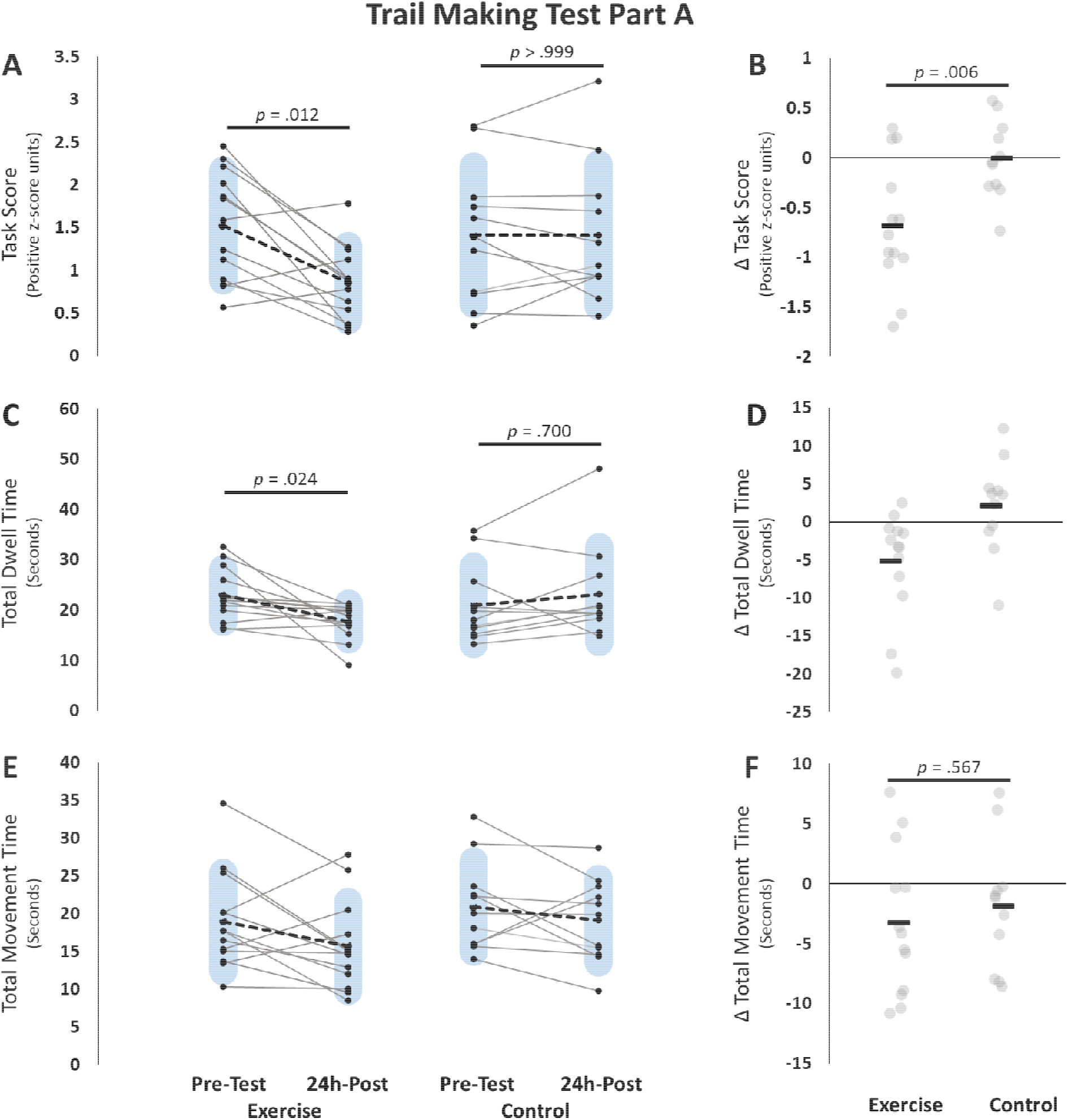
Trail Making Test Part A (TMT-A). Pre-testing and 24h-post testing participant values for the Exercise group (left) and control group (right) for **A)** Task score, **C)** Total dwell time, **E)** Total movement time. Panels **B), D)**, and **F)** represent change scores for each of the measures respectively. group × time RM-ANOVA Interaction results are represented with the line and p-values over the change scores on the right. Total dwell time was run with non-parametric testing and therefore no group × time interaction was assessed. All p-values are Bonferroni adjusted. Blue shaded bars represent standard deviation. For panels B, D, and F Black bars represent the group means. Grey circles represent individual data points

Since an effect was only observed for the exercise group, only it was used to investigate sex differences with a sex (male, female) × time RM-ANOVA. For this analysis, only the sex × time interaction and the main effect of group were reported given the main effect of time is redundant with what has already been reported in the previous analyses. The RM-ANOVA test did not detect a sex × time interaction [*F*(1, 11) = .104, *p* = .753, □_p_^2^= .009], or a main effect of group [*F*(1, 11) = .052, *p* = .824, □_p_^2^= .005].

#### Dwell time

For TMT-A dwell time, there was a homogeneity of variance violation (24h-post, *p* = .05), and therefore the higher order RM-ANOVA was not assessed. Mann-Whitney U tests for between group comparisons at pre-testing (*W* = 48, *p* = .744, Rank-Biserial Correlation = −.329) and 24h-post (*W* = 93, *p* = .912, Rank-Biserial Correlation = .301) were not significant. A Wilcoxon signed-rank test revealed a significant decrease in dwell time for the exercise group (*W* = 83, *z* = 2.621, *p* = .024, Rank-Biserial Correlation = .824), but not for the control group (*W* = 17, *z* = −1.423, *p* = .700, Rank-Biserial Correlation = −.485; Figure 4).

Once again, only an effect was observed for the exercise group, and therefore only the exercise group data were examined for sex differences. Separate Mann-Whitney U tests at pre-testing (*W* = 16, *p* > .999, Rank-Biserial Correlation = .067) and 24h-post (*W* = 17, *p* > .999, Rank-Biserial Correlation = .133) failed to detect any differences between sexes. Although separate Wilcoxon signed rank tests for males (*W* = 53, *z* = 2.599, *p* = .024, Rank-Biserial Correlation = .927) and females (*W* = 4.00, *z* = .535, *p* > .999, Rank-Biserial Correlation = .333) revealed that only males improved their processing speed after exercise.

#### Movement time

For TMT-A, a RM-ANOVA revealed a significant main effect of time [*F*(1, 22) = 4.629, *p* = .043, □_p_^2^= .174] indicating that both groups decreased their movement speed over the course of the intervention. However, the group × time interaction [*F*(1, 22) = .339, *p* = .567, □_p_^2^= .015] and the main effect of group [*F*(1, 22) = 1.563, *p* = .224, □_p_^2^= .066] were not statistically significant. These findings suggest that there were no differences between groups in change in movement speed after five-days of motor training (Figure 4).

### Trail making test part B

#### Task score

For TMT-B Task score, normality (*W* = .871, *p* = .005), and homogeneity of variance (24h-post, *p* = .004) were violated. Therefore, non-parametric Mann-Whitney U tests were used to determine there were no between group differences at pre-testing (*W* = 95.00, *p* = .744, Rank-Biserial Correlation = .329), and 24h-post (*W* = 78.00, *p* > .999, Rank-Biserial Correlation = .091) timepoints. Separate Wilcoxon signed-rank tests were used to determine that there were also no within group differences between pre-testing and 24h-post testing for the exercise group (*W* = 33.00, *z* = −.874, *p* > .999, Rank-Biserial Correlation = −.275) or the control group (*W* = 52.00, *z* = 1.689, *p* = .408, Rank-Biserial Correlation = .576; Figure 5).

**Figure 5.**
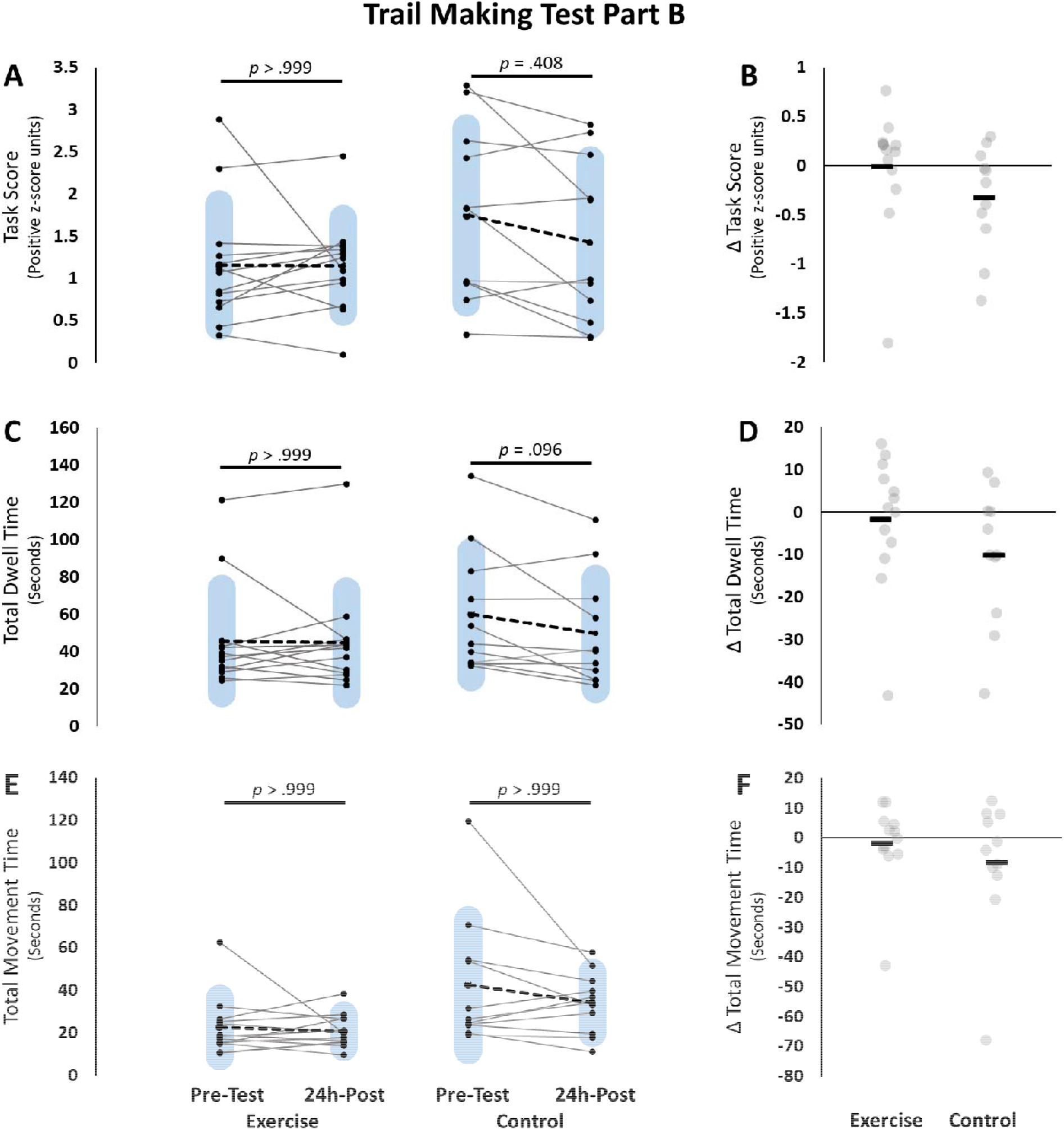
Trail Making Test Part B (TMT-B). Pre-testing and 24h-post testing participant values for the Exercise group (left) and control group (right) for **A)** Task score, **C)** Total dwell time, **E)** Total movement time. Panels **B), D)**, and **F)** represent change scores for each of the measures respectively. All tests were run with non-parametric testing and therefore no group × time interactions were assessed. All p-values are Bonferroni adjusted. Blue shaded bars represent standard deviation. For panels B, D, and F Black bars represent the group means. Grey circles represent individual data points

#### Dwell time

For TMT-B, the change values from pre-testing to 24h-post were not normally distributed based on Shapiro-Wilk’s test of normality (*W* = .905, *p* = .027). Therefore, non-parametric Mann-Whitney U tests were performed on pre- and 24h-post testing time points between groups, and these analyses failed to detect differences at pre-testing (*W* = 97.00, *p* = .600, Rank-Biserial Correlation = .357), and 24h-post (*W* = 63.00, *p* > .999, Rank-Biserial Correlation = −.119). Finally, separate Wilcoxon signed-rank tests suggest that both the exercise (*W* = 33.00, *z* = −.874, *p* > .999, Rank-Biserial Correlation = −.275) and control (*W* = 58.00, *z* = 2.223, *p* = .096, Rank-Biserial Correlation = .758) groups did not reduce their dwell time from pre-testing to 24h-post timepoints (Figure 5).

#### Movement time

For TMT-B, the change values from pre-testing to 24h-post were not normally distributed based on Shapiro-Wilk’s test of normality (*W* = .762, *p* = .003), and Levene’s homogeneity of variance tests were violated at pre-testing (*p* = .019) and 24h-post (*p* = .050). Therefore, non-parametric Mann-Whitney U tests were carried out to assess between group differences at pre-testing (*W* = 107.00, *p* = .164, Rank-Biserial Correlation = .497), and 24h-post (*W* = 111.00, *p* = .088, Rank-Biserial Correlation = .552) timepoints. These analyses did not show any differences between groups in TMT-B movement time. Additionally, no differences between time points were observed for either the exercise group (*W* = 49.00, z *=* .245, *p* > .999, Rank-Biserial Correlation = .077), or the control group (*W* = 44.00, z *=* .*978, p* > .999, Rank-Biserial Correlation = .333; Figure 5).

### Resting-state functional connectivity

Since different MRI scanners were used in this study (Philips Achieva: n = 14; exercise = 12; control = 2; Philips Elition: n = 11; exercise = 2; control = 9) and there were clear differences between groups for which scanner was used, we first assessed whether scanner type was a significant covariate in a group × time RM-ANOVA model for DLPFC-sensorimotor network functional connectivity. Scanner type was not a significant covariate [*F*(1,22)= .197, *p* = .661, □_p_^2^= .009] and therefore no adjustments to the statistical models were made. A group-level seed-to-network functional connectivity analysis revealed a decrease in functional connectivity between the right DLPFC (seed) and the sensorimotor network in the exercise group only (family-wise error rate corrected *p* = .024). Specifically, within the sensorimotor network, a significant cluster of decreased functional connectivity was observed over the left inferior parietal lobule [IPL, 232 voxels, CoG MNI coordinates: X = −58.91, Y = −25.05, Z = 20.58; Figure 6]. We did observe a significant pre-testing difference between groups in the DLPFC-sensorimotor network functional connectivity [*t*(23) = −4.589, *p* < .001, Cohen’s *d* = −1.849]. To determine if this pre-testing difference impacted our results, an additional between groups analysis of covariance test was run with pre-testing DLPFC-sensorimotor network functional connectivity serving as the covariate. This analysis revealed that a significant difference remained between groups (exercise: −5.841 ± 10.844; control: −2.817 ± 8.474) after covarying pre-testing differences [*F*(1,22) = 5.623, *p* = .027, □_p_^2^= .204]. To visualize the participant-level change in functional connectivity that gave rise to this significant cluster, participant-level connectivity values at each time point were extracted using *fslmeants*. These functional connectivity values were then used in correlation analyses to determine if the change in functional connectivity was related to the changes in TMT-A performance.

**Figure.**
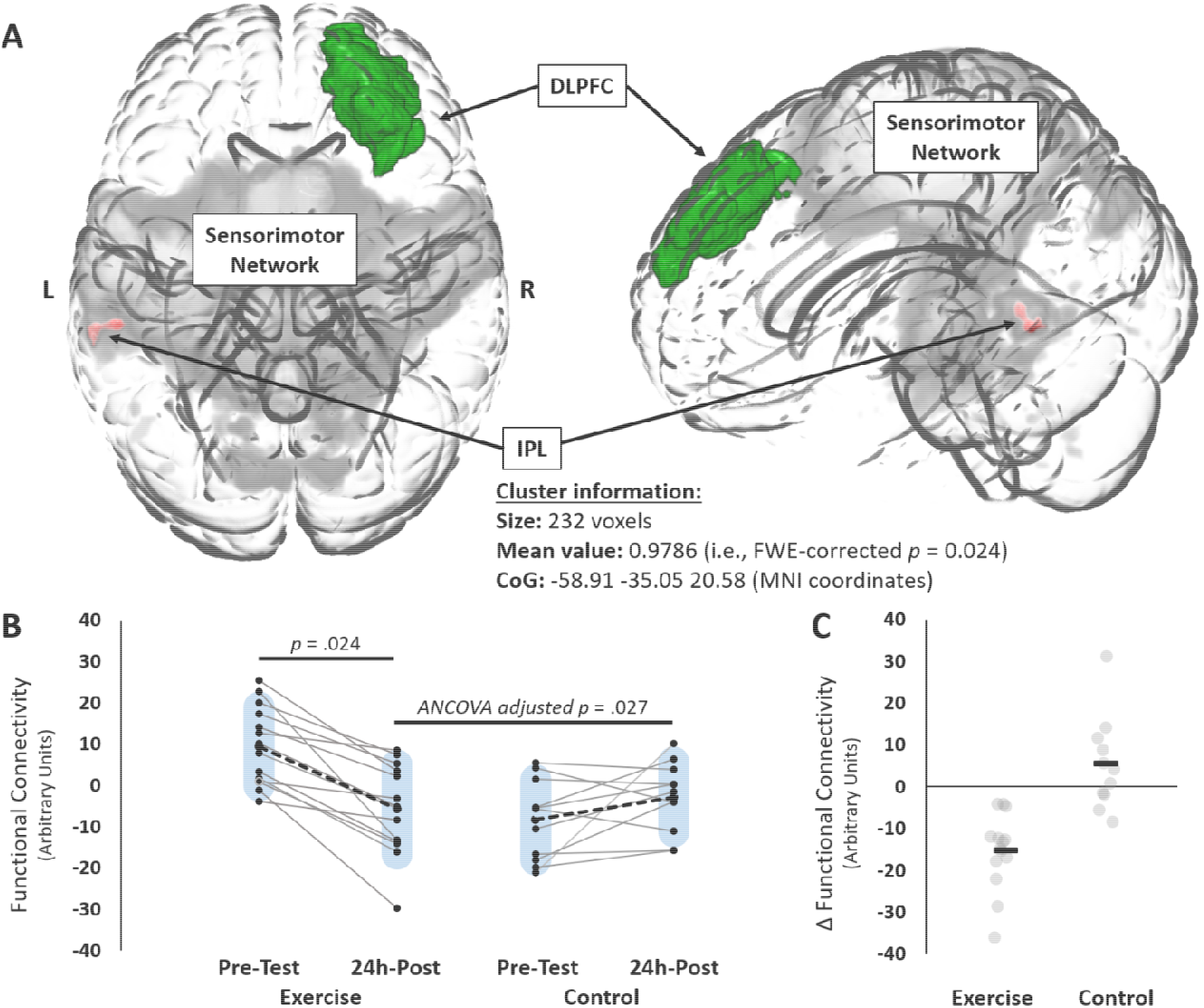
Error! No text of specified style in document.6. Resting-state functional connectivity results from the seed-to-network analysis. **A)** A glass brain showing the right dorsolateral prefrontal cortex (DLPFC) seed in green, the sensorimotor network component from the group level ICA (grey), and the significant cluster of decreased functional connectivity over the left inferior parietal lobule (IPL) in red for the exercise group. **B)** individual functional connectivity scores at pre-testing and 24h-post time points for the exercise group (left) and the control group (right). **C)** Functional connectivity change scores (24h-post – pre-testing) for the exercise group (left) and control group (right). Blue shaded bars represent standard deviation. For panel C, black bars represent the group means. Grey circles represent individual data points ANCOVA = Analysis of Covariance adjusted for pre test differences.

To determine if the significant decrease in DLPFC-sensorimotor network functional connectivity was a by-product of global changes in functional connectivity, we investigated the intra-network functional connectivity of the sensorimotor network. This analysis did not show any statistically significant differences between groups or time points.

Since an effect was only observed for the exercise group, only it was used to investigate sex differences with a sex (male, female) × time RM-ANOVA. For this analysis, only the sex × time interaction and the main effect of group were examined given the main effect of time is redundant with what has already been reported in the previous analyses. The RM-ANOVA failed to detect a significant sex × time interaction [*F*(1, 12) = 1.480, *p* = .247, □_p_^2^= .110], or a main effect of sex [*F*(1, 12) = 1.121, *p* = .311, □_p_^2^= .085].

### Relationship between functional connectivity and motor learning

To determine if motor learning of the serial targeting task correlated with the change in DLPFC-sensorimotor network functional connectivity, a Spearman’s Rho correlation was performed. The Spearman’s rho correlation failed to detect a significant relationship between these two variables (ρ = .389, *p* = .067), indicating that the improvement in motor learning was not related to the change in functional connectivity between the DLPFC and the sensorimotor network.

### Relationship between functional connectivity and improved processing speed

After observing significant effects for TMT-A task score and dwell time, in addition to a significant cluster of decreased functional connectivity between the DLPFC-sensorimotor network for the exercise group, we sought to determine if a change in functional connectivity was related to a change in cognitive-motor performance for both groups. With both groups data pooled together, Spearman’s rho correlations between the change in functional connectivity and the change in TMT-A task score (ρ = .458, *p* = .025) and dwell time (ρ = .418, *p* = .043) were both positively correlated. These relationships suggest that the individuals that experienced a greater reduction in DLPFC-sensorimotor network functional connectivity also improved their overall task performance, and reduced the time needed to visually scan for the target and plan their next arm movement (Boyd *et al*., 2009*b*) (Figure 7).

**Figure 7.**
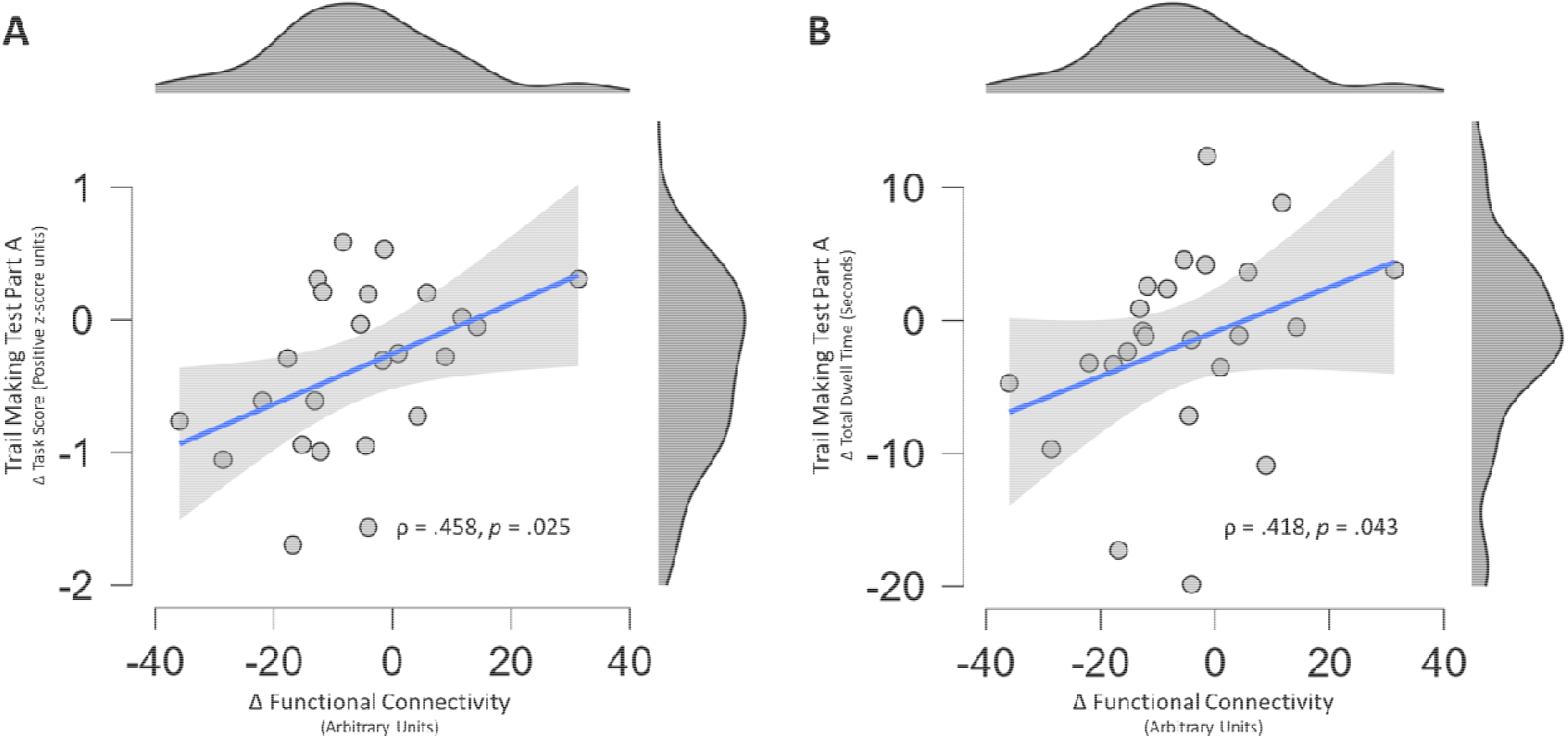
Spearman’s Rho (ρ) correlations between the change in DLPFC-sensorimotor network functional connectivity (x-axes), and the change in Trail Making Test Part A (TMT-A) **A)** Task score, and **B)** Total dwell time. The Grey shaded areas represent the 95% confidence interval around the regression line.

## Discussion

In the present study, we investigated the impact of high intensity aerobic exercise paired with motor training on cognitive-motor function in individuals living with chronic stroke. After a five-day intervention pairing either exercise or rest before a paretic arm implicit motor learning task, we observed significant improvements in cognitive processing speed with TMT-A, but not with TMT-B with the less-affected arm. We also used resting-state functional brain imaging to determine if changes in functional connectivity between the DLPFC and the sensorimotor network would be observed after our intervention and whether these changes would relate to behavioural changes. We observed a decrease in DLPFC-sensorimotor network functional connectivity that was correlated with a change in overall TMT-A task performance (ρ = .453), and the TMT-A dwell time (ρ = .418) regardless of group.

### Cognitive-motor performance is enhanced after exercise

In the present study, we saw an improvement in TMT-A performance for the exercise group but not the control group, using the TMT-A task score. After separating the cognitive and motor components of TMT-A, it was evident that pairing high intensity aerobic exercise with motor training had a positive impact on performance in the cognitive domain, this was supported by a significant decrease in dwell time for the exercise group only, with no between group differences for movement time.

The decreased dwell time for TMT-A and not TMT-B hints at how exercise differentially impacts the neurocognitive processes involved in these tasks. TMT-A is a measure of cognitive-motor processing speed (Bowie & Harvey, 2006), whereas TMT-B assesses more complex cognitive processes like task switching and mental flexibility (Bowie & Harvey, 2006). Our data suggest that high intensity aerobic exercise had a specific impact on processing speed rather than on complex cognitive processes such as task switching and mental flexibility.

### Altered resting-state functional connectivity related to cognitive-motor performance

In the present study, an expected decrease in functional connectivity was observed between the right DLPFC, which is known for its involvement in processing speed (Hillary *et al*., 2006; Kaller *et al*., 2011) and response selection (Boyd *et al*., 2009*b*), and the sensorimotor network in the exercise group only. Specifically, the cluster of decreased functional connectivity within the sensorimotor network was found over the left inferior parietal lobule, which is involved in action planning and prediction (Wolpert & Ghahramani, 2000; Kilner *et al*., 2007; Elk, 2014). The decrease in functional connectivity between these regions after exercise may reflect a beneficial effect of exercise on cognitive-motor processing speed in individuals with stroke, whereby the decreased coupling reflects a shift towards automaticity of perception and action, and illustrates a reduced dependence on cognitive resources to complete a cognitively demanding motor task (Mazzoni, 2008; Wu *et al*., 2008). This notion is supported by previous research that found a decrease in DLPFC BOLD signal after learning a cognitively challenging repeated sequence continuous target tracking task with healthy controls, but not in individuals with stroke (Meehan *et al*., 2011). These findings likely coincide with “slow” or “late” phases of motor learning (Dayan & Cohen, 2011). In these stages of learning the attentional demand and executive resources are no longer required for effective task execution (Schneider & Shiffrin, 1977; Doyon & Benali, 2005; Ashby *et al*., 2010; Wu *et al*., 2015). However, past work suggested that motor practice alone was insufficient to stimulate automaticity of motor plans after stroke; importantly in this previous work, the same dose of practice enabled age matched healthy controls to reduce their reliance on DLPFC suggesting that they automated learned movements (Meehan *et al*., 2011). Critically, the current study suggests that motor training paired with high intensity aerobic exercise facilitates the acquisition of TMT-A after stroke, which is a cognitive-motor task that specifically relies on processing speed.

In the present study, we observed significant motor learning improvements in both groups (one sample Wilcoxon signed-rank test: *p* < .001). However, there were no differences between groups (*p* = .738), and the improvements were not related to the change in functional connectivity. Previous research from our lab using the same experimental paradigm in a healthy aging cohort (Greeley *et al*., 2021) and individuals with chronic stroke (Greeley *et al*., 2023) also failed to see a preferential advantage of high intensity aerobic exercise for enhancing implicit motor sequence learning on the serial targeting task compared to controls. These findings may suggest implicit motor sequence learning tasks, which do not rely heavily on the prefrontal cortex, do not need the benefits conferred by acute bouts of high intensity aerobic exercise to be learned; instead, they potentially rely on plasticity within motor networks. In contrast, our data show that tasks that require cognitive-motor interactions appear to benefit greatly from an intervention that amplifies plasticity in the prefrontal cortex.

## Limitations and future directions

In the context of the present study, high intensity aerobic exercise paired with motor training failed to alter cognitive-motor performance in TMT-B, which depends on mental flexibility and task switching. Future work may explore alternative manipulations to various exercise variables such as duration or intensity of exercise bouts, frequency of exercise sessions, the timing of exercise sessions in proximity to motor training or even explore anaerobic or resistance exercise training modalities to determine their efficacy for improving not only processing speed but other more complex neuro-cognitive processes. It is also currently unclear how long the exercise-related effects on TMT-A would be retained, and future work should consider investigating this phenomenon with an additional delayed retention test after the intervention. In addition, there was no differential impact of high intensity aerobic exercise on motor learning as characterized by B scores. Importantly this shows that for an implicit motor sequence learning task (the serial targeting task) practice alone enabled both groups to learn. Future work should consider more complex, cognitive-motor learning tasks to understand what types of skills benefit from being paired with high intensity aerobic exercise. We also paired exercise with motor practice in this study, and it is unclear whether exercise alone would produce similar outcomes. Therefore, future research should explore the effects of exercise on cognitive-motor processing speed without skilled motor practice. Finally, the participant sample in the exercise group was predominantly male (n =19) compared to female (n = 6). This biological sex imbalance limits our ability to accurately interpret sex differences and future work should consider larger sample sizes with a more balanced sex distribution to be able to adequately explore if our findings translate equally or differ between sex.

## Conclusions

Five-sessions of high intensity aerobic exercise paired with skilled motor training improved cognitive-motor performance on a processing speed dependent task. Interestingly, this effect was not observed for a more complex cognitive-motor task that depended on task switching and mental flexibility. We also observed a relationship between the amount of change in DLPFC-sensorimotor network functional connectivity and the change in overall task performance, and processing speed during TMT-A. Regardless of group, the individuals that had greater reductions in functional connectivity performed better on TMT-A. These findings suggest that in individuals with chronic stroke, high intensity aerobic exercise may lead to brain changes that enable a beneficial decrease in cognitive resources dedicated to task execution, thereby correcting the high cognitive demand of complex motor tasks often seen after stroke (Puh *et al*., 2007; Li *et al*., 2014; Hall *et al*., 2021). This intervention allowed a restoration of cognitive-motor function that may have meaningful effects on complex motor task performance in individuals with chronic stroke.

## Acknowledgements

The authors would like to acknowledge Daniela J. Palombo Ph.D. from the Department of Psychology at the University of British Columbia for her valuable insight and feedback with analyses, and interpretation of this work.

